# *Mycobacterium ulcerans* low infectious dose and atypical mechanical transmission support insect bites and puncturing injuries in the spread of Buruli ulcer

**DOI:** 10.1101/071753

**Authors:** John R. Wallace, Kirstie M. Mangas, Jessica L. Porter, Renee Marcsisin, Sacha J. Pidot, Brian Howden, Till F. Omansen, Weiguang Zeng, Jason K. Axford, Paul D. R. Johnson, Timothy P. Stinear

## Abstract

Addressing the transmission enigma of the neglected disease Buruli ulcer (BU) is a World Health Organization priority. In Australia, we have observed an association between mosquitoes harboring the causative agent, *Mycobacterium ulcerans*, and BU. Here we tested a contaminated skin model of BU transmission by dipping the tails from healthy mice in cultures of the causative agent, *Mycobacterium ulcerans*. Tails were exposed to mosquito (*Aedes notoscriptus* and *Aedes aegypti*) blood feeding or punctured with sterile needles. Two of 11 of mice with *M. ulcerans* contaminated tails exposed to feeding *A. notoscriptus* mosquitoes developed BU. Eighteen of 20 mice subjected to contaminated tail needle puncture developed BU. Mouse tails coated only in bacteria did not develop disease. We observed a low infectious dose-50 of four colony-forming units and a median incubation time of 12 weeks, consistent with data from human infections. We have uncovered a highly efficient and biologically plausible atypical transmission mode of BU via natural or anthropogenic skin punctures.

**Author summary:** Buruli ulcer is a neglected tropical disease caused by infection with *Mycobacterium ulcerans*. Unfortunately, how people contract this disease is not well understood. Here we show for the first time using experimental infections in mice that a very low dose of *M. ulcerans* delivered beneath the skin by a minor injury caused by a blood-feeding insect (mosquito) or a needle puncture is sufficient to cause Buruli ulcer. This research provides important laboratory evidence to advance our understanding of Buruli ulcer disease transmission.

## Introduction

Among the 17 neglected tropical diseases the World Health Organization (WHO) has targeted for control and elimination, only Leprosy and Buruli ulcer (BU) have unknown modes of transmission [1]. The search to understand how humans contract BU spans more than 70 years since the causative agent, *Mycobacterium ulcerans*, was first identified [2]. There are persistent and emerging foci of BU cases across the world, in particular Africa and Australia [3]. BU is characterized by necrotizing skin lesions, caused by localized proliferation of *M. ulcerans* in subcutaneous tissue. BU is rarely fatal, but untreated infections leave patients with significant disfigurement and disability, with damaging personal and economic consequences [4, 5]. Researchers have long been struck by the characteristic epidemiology of BU, with cases occurring in highly geographically circumscribed regions (sometimes less than a few square kilometres) and risk factors for infection that include gardening, insect bites and proximity to (but not necessarily contact with) lacustrine/riverine regions [6–14]. Human-to-human spread is considered unlikely [14]. Disease transmission is thought to occur by contact with an environment contaminated with *Mycobacterium ulcerans* but exactly where the pathogen resides and why it appears so geographically restricted have yet to be determined. [15].

*M. ulcerans* is very slow growing (doubling time >48 hrs) and this poses a problem for source tracking efforts as it is difficult to isolate the bacteria in pure culture from complex environmental specimens [16]. *M. ulcerans* has only once been isolated from a non-clinical source, an aquatic water bug (Gerridae) from Benin [16]. Quantitative PCR targeting *M. ulcerans*-specific DNA is the most frequently used technique in surveys of environmental specimens. A comprehensive review of the many field and lab studies that have examined reservoir and transmission of BU has highlighted the range of organisms from aquatic insects, fish, amphibia, and in Australia certain native marsupials that can serve as potential reservoirs for *M. ulcerans* [15, 17]. Since the first observation that biting aquatic insects can harbour *M. ulcerans* [18], studies of BU transmission have largely focused on the potential for insects to biologically vector *M. ulcerans* implying that *M. ulcerans* undergoes a propagative or reproductive mode of development in an insect [19–23]. Several case-control studies, including from both Australia and Africa have suggested insects may play a role in transmission [10, 11]. However, there is no compelling experimental evidence for single-mode biological transmission of *M. ulcerans* via insect vectors.

In southeastern Australia, we noted Buruli lesions on exposed areas likely to attract biting insects, some patients with every brief exposure times to endemic areas [24, 25] and 2004 we began a study that identified *M. ulcerans* DNA associated with mosquitoes captured in endemic areas [19].

## Materials and Methods

### Bacterial isolates and culture conditions

*M. ulcerans* strain JKD8049 and bioluminescent *M. ulcerans* JKD8049 (harbouring plasmid pMV306 *hsp:luxG13*) [26, 27] were cultured in 7H9 broth or Middlebrook 7H10 agar, containing 10% oleic-albumin-dextrose-catalase growth supplement (Middlebrook, Becton Dickinson, Sparks, MD, USA) and 0.5% glycerol (v/v) at 30°C. Colony counts from bacterial cultures or tissue specimens were performed using spot plating. Five x 3μl volumes of serial 10-fold dilutions (10^−1^ to 10^−5^) of a culture or tissue preparation were spotted onto 7H10 agar plates with a 5x5 grid marked. The spots were allowed to dry, the plates loosely wrapped in plastic bags and then incubated as above for 10 weeks before counting colonies. Data analysis was performed using GraphPad Prism (v 6.0). All culture extracts were screened by LC-MS for the presence of mycolactones as previously described to ensure bacteria used in transmission experiments remained fully virulent [28].

### Experimental animals

The animal ethics committee (AEC) of the University of Melbourne approved all animal experiments under approval number AEC: 1312775.2. DBALB/c mice were purchased from ARC (Canning Vale, Australia) and housed in individual ventilated cages. Upon arrival, animals were acclimatizing for 5 days. Food and water were given *ad libitum*.

*Aedes notoscriptus and Aedes aegypti rearing*. Wild caught mosquitoes were sourced from around Cairns, Queensland, Australia. *A. notoscriptus* and *A. aegypti* colonies were reared in a Physical Containment Level 2 (PC2) laboratory environment at 26 °C using previously described methods, with the addition of brown paper used as the oviposition substrate for *A. notoscriptus* [29].

### Mosquito-mouse transmission experiments

Four-week old female BALB/c mice were anaesthetized and their tails coated in a thin film of *M. ulcerans* by dipping the tails in a Petri dish containing 20mL of bacterial culture (concentration ~10^6^ CFU/mL). The tail only was then exposed to a 200mm x 200mm x 200mm cage of 20 adult female mosquitoes for a period of 15 minutes. The number of insects biting each mouse was recorded over the exposure period by continuous observation. Mice were then observed weekly for up to six months for signs of tail lesions. Sterile needle stick (25G or 30G needle) and no-trauma were used as controls. An additional control consisted of tails dipped in sterile culture broth only and subjected to mosquito biting or sterile needle stick.

*Real time quantitative PCR*. For each mosquito that blood-fed DNA was individually extracted from the dissected head, abdomen and legs of each insect using the Mo Bio Powersoil DNA extraction kit following manufacturer’s instructions (Mo Bio Laboratories Inc., Carlsbad CA USA). DNA was similarly extracted from mouse tissue. Procedural extraction control blanks (sterile water) were included at a frequency of 10% to monitor potential PCR contamination, in addition to no-template negative controls. *IS2404* quantitative PCR (qPCR) was performed as described [30]. IS2404 cycle threshold (Ct) values were converted to genome equivalents (GE) to estimate bacterial load within a sample by reference to a standard curve (r^2^=0.9312, y=[-3.000Ln(x)+39.33]*Z, where y=Ct and x=amount of DNA [fg] and Z=the dilution factor]), calculated using dilutions of genomic DNA from *M. ulcerans* strain JKD8049, quantified using fluorimetry (Qubit, Invitrogen) [30].

*Preparation of mouse tissue for analysis*. At the end of the experimental period or when a clinical end-point was reached mice were humanely killed. The region of a mouse-tail spanning a likely lesion was cut into three equal sections for histology, qPCR and CFU counts. Individual tail pieces for CFU counts were weighed and placed into sterile 2ml screw capped tube containing 0.5g of large glass beads and 600pl of sterile 1x PBS. Tissues were homogenized using four rounds of 2x 30second pulses in a high-speed tissue-disruptor at 6500 rpm, with tubes placed on ice for 5 minutes between each round. A 300μl volume of this homogenate was decontaminated with 300μl of 2% NaOH (v/v) and incubated at room temperature for 15 minutes. The preparation was neutralized drop-wise with a 10% solution of orthophosphoric acid (v/v) with added bromophenol blue until the solution changed from blue to clear. The mixtures were diluted in PBS and CFUs determined by spot plating as described above.

*Histology*. Sections of mouse-tails were fixed in 10% (w/v) neutral-buffered-formalin and imbedded in paraffin. Each mouse-tail was sectioned transversely (four micron thickness) and subjected to Ziehl-Neelson and hematoxylin/eosin staining. The fixed and stained tissue sections were examined by light microscopy.

*Infectious Dose*. To estimate the infectious dose we measured the surface area of five dissected mouse-tails to obtain an average surface area (493.3 ± 41.1 mm^2^). Using ten mouse-tails, we then calculated the average volume of *M. ulcerans* 7H9 Middlebrook culture adhering to the tail surface (32.4 ± 4.2 mL), the concentration of bacteria in the cultures used, and the surface area of the tips of 25G and 30G needles used to deliver the puncture wounds (0.207 mm^2^ and 0.056 mm^2^ respectively). These parameters were then used to calculate the infectious dose, assuming the bacteria were evenly distributed over the tail surface (Fig. 1C). A standard curve was interpolated using non-linear regression and an ID_50_ estimated using (GraphPad Prism v 7.0a).

## Results

### M. ulcerans is efficiently transmitted to a mammalian host by an atypical mechanical means

We established a murine model of *M. ulcerans* transmission (model-1) that represented a skin surface contaminated with the bacteria and then subjected to a minor penetrating trauma, via either a mosquito bite or needle stick puncture. In a first experiment two of six mice with their tails coated with *M. ulcerans* and then bitten by mosquitoes developed lesions (Table 1, Fig. 1C, Fig. 2A). Histology of these lesions confirmed a subcutaneous focus of AFB, within a zone of necrotic tissue. There was also characteristic epithelial hyperplasia adjacent to the site of infection (Fig. 2B,C). Material extracted from the lesions was *IS2404* qPCR-positive and culture positive for *M. ulcerans* (Supplementary Table S1). Mice bitten by mosquitoes but with tails coated only with sterile culture media did not develop lesions (Table 1). In the same experiment, we also subjected five mice to a single needle stick puncture. Each mouse had their tail coated with *M. ulcerans* as for the mosquito biting. Four of these five mice developed *M. ulcerans* positive lesions (Table 1, Fig. 2D), with subcutaneous foci of infection and viable bacteria (Fig. 2F). Six mice with their tails coated with *M. ulcerans* but not subjected to a puncturing injury did not develop lesions and remained healthy until the completion of the experiment at six months. This experiment suggested that minor penetrating skin trauma (defined here as a puncture <0.5mm diameter and <2mm deep) to a skin surface contaminated with *M. ulcerans* is sufficient to cause infection. It also revealed a means by which mosquitoes could act as atypical mechanical vectors of *M. ulcerans*.

**Table 1:** Summary of transmission experiments.

| Trauma source | Number of mice | Mouse tail coated | Number of mice bitten | Number of mice developing BU | Estimated dose (CFU) |
| --- | --- | --- | --- | --- | --- |
| **Experiment 1:**<br>(Atypical mechanical transmission model, contaminated skin surface) |  |  |  |  |  |
| *Aedes notoscriptus* | 12<br>(4/mosquito cage)* | *M. ulcerans*<br>(4.1x 10<sup>6</sup> CFU/mL) | 6 | 2<br>(#182, #191) | 21 |
| *Aedes notoscriptus* | 4<br>(1/mosquito cage) | Media-only | 2 | 0 | - |
| Sterile needle (25G) | 5 | *M. ulcerans*<br>(4.1x 10<sup>6</sup> CFU/mL) | - | 4<br>(#186, #200, #201, #202) | 55 |
| None | 6 | *M. ulcerans*<br>(4.1x 10<sup>6</sup> CFU/mL) | - | 0 | - |
| **Experiment 2:**<br>(Atypical mechanical transmission model, contaminated skin surface) |  |  |  |  |  |
| *Aedes aegypti* | 5<br>(1/mosquito cage) | *M. ulcerans*<br>(1.83 x 10<sup>6</sup> CFU/mL) | 5<sup>#</sup> | 0 | 9 |
| *Aedes aegypti* | 3<br>(1/mosquito cage) | Media-only | 2 | 0 | - |
| Sterile needle (25G) | 5 | *M. ulcerans*<br>(1.83 x 10<sup>6</sup> CFU/mL) | - | 4<br>(#216, #217, #218, #219) | 40 |
| None | 5 | *M. ulcerans*<br>(1.83 x 10<sup>6</sup> CFU/mL) | - | 0 | - |

**Notes:** \*20 adult female mosquitoes per cage; <sup>#</sup>Multiple bites per mouse with 2 mice receiving 3 bites and 1 mouse receiving 2 bites; <sup>§</sup>“GE” is *M. ulcerans* genome equivalents as estimate based on IS2404 qPCR.

**Fig. 1:**
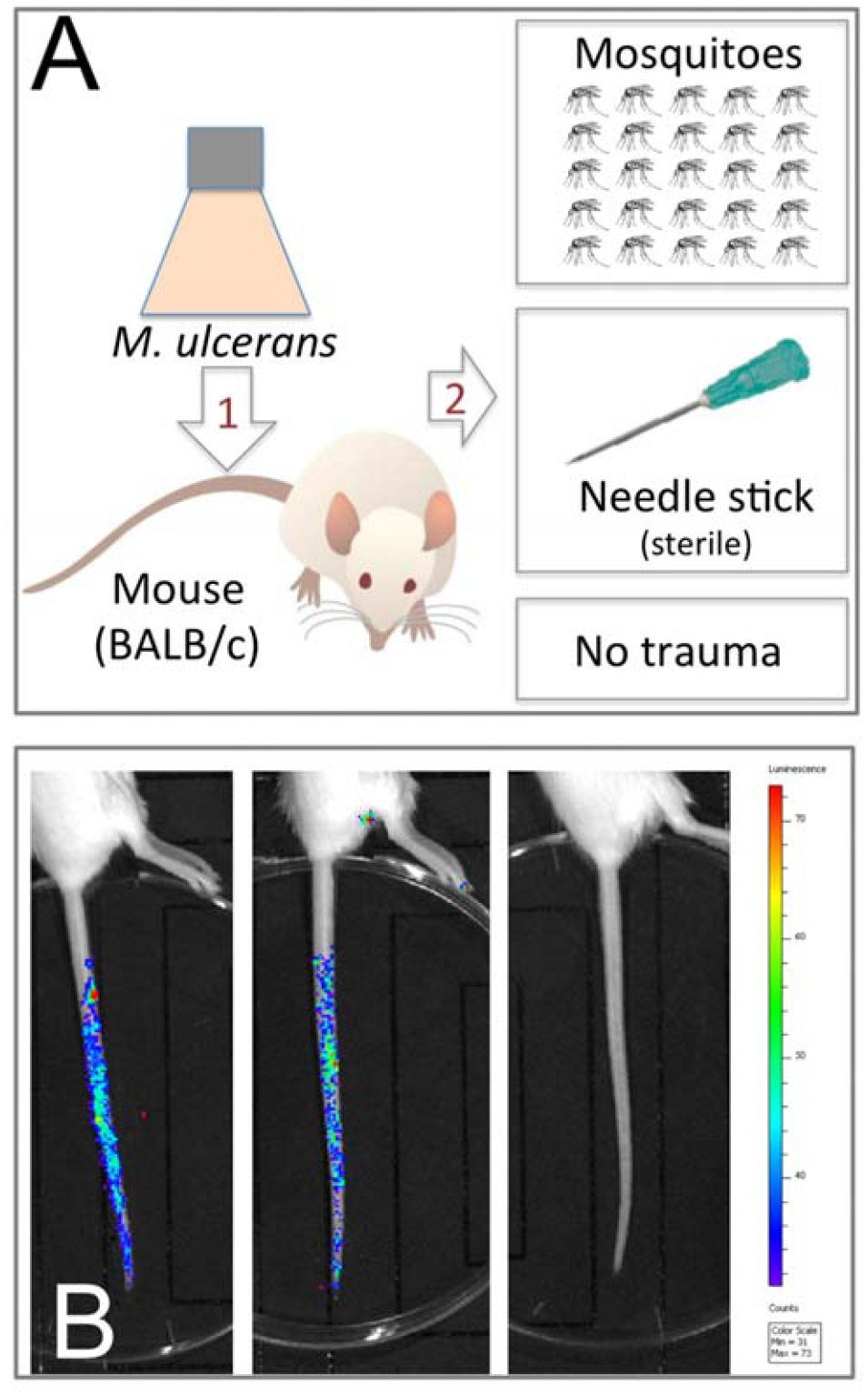
**Schematic representations of the two BU transmission models tested in this study.** (A) Model-1 tests transmission of *M. ulcerans* present on a skin surface following a puncturing injury created by mosquito blood-feeding or needle stick. (B) Visualization of bioluminescent *M. ulcerans* JKD8049 (harbouring plasmid pMV306 *hsp:luxG13*) [26, 27] on the mouse-tail in model-1, showing the distribution of bacteria immediately after coating for two mice, versus an uncoated animal. *M. ulcerans* concentration was 10^6^ CFU/mL.

**Fig. 2:**
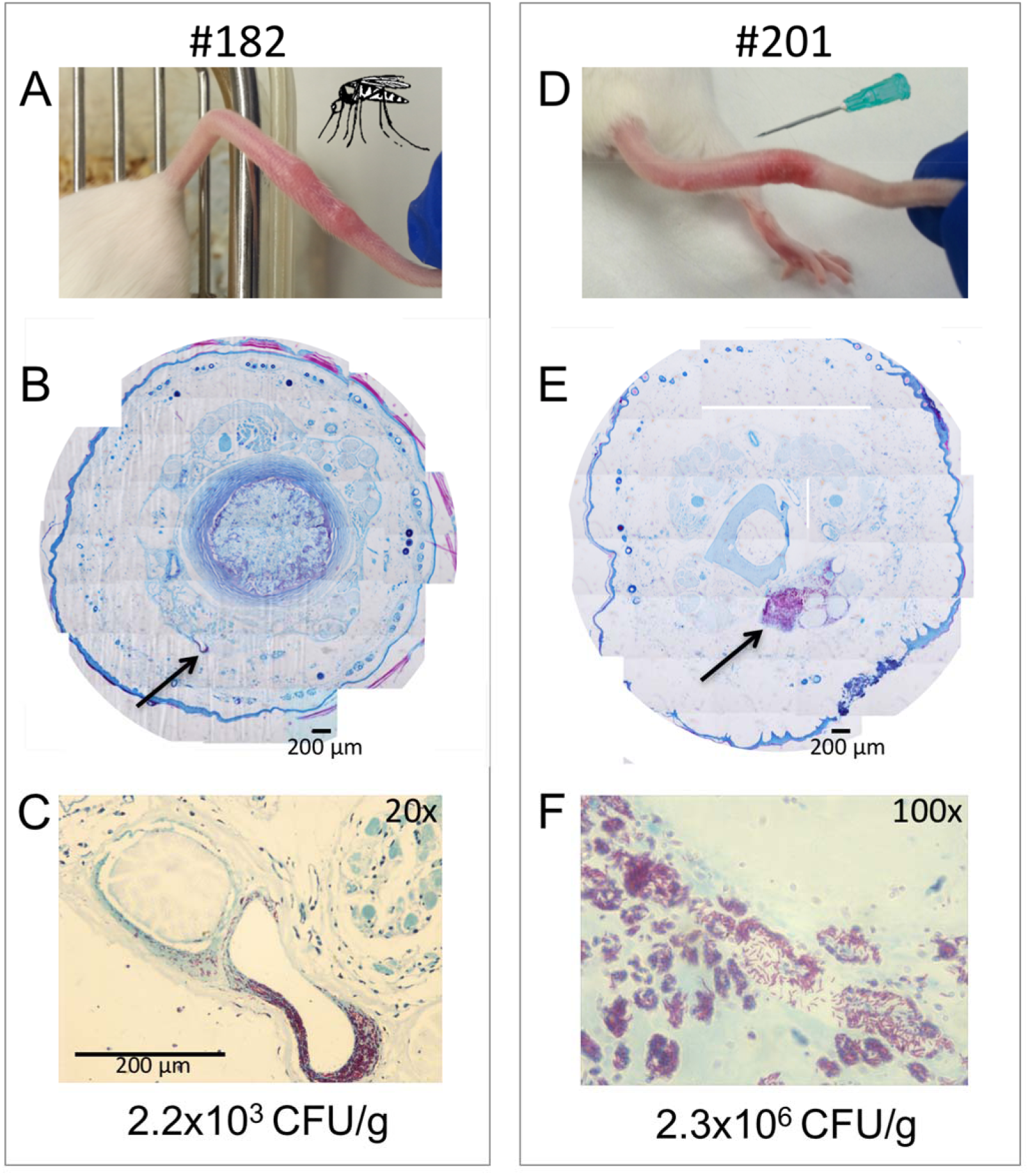
**Atypical mechanical transmission of *M. ulcerans***. (A) An example of the development of Buruli ulcer following mosquito blood-feeding through a skin surface (mouse-tail) contaminated with *M. ulcerans.* (B) Composite histological cross-section with Ziehl-Neelsen staining through the infected tail showing the focus of AFB bacteria (arrow) within the subcutaneous tissue. (C) Higher magnification view of the focus of infection, with the yield of viable M. ulcerans obtained from the infected tissue. Panels (D) − (F) show the same analyses as for the mosquito-bitten mouse #182, but for a mouse developing a lesion following sterile needle-stick puncture through a contaminated skin surface (mouse #201).

### M. ulcerans burden on mosquitoes correlates with transmission

Then, using approximately the same dose of bacteria to coat the mouse-tails, we repeated experiment-1 but with *Aedes aegypti* because of the close association of this mosquito to humans world-wide and their vector competency for viral pathogens. Despite more mosquito bites per mouse than the first experiment, none of the five insect-exposed mice developed lesions (Table 1). In contrast however, four of five mice subjected to single, needle stick puncture developed *M. ulcerans* positive tail lesions (Table 1). We assessed the burden of *M. ulcerans* by individual IS2404 qPCR of the head, abdomen and legs for each mosquito that blood fed (Fig. 3). A summary of these results is shown in Fig. 3A. We noted that the bacterial load (expressed as genome equivalents [GE]) was significantly higher in the heads of mosquitoes associated with mice that developed lesions (p<0.05) (Fig. 3B, Supplementary Table S1). These data point to a threshold, above which mosquitoes can become competent mechanical vectors for *M. ulcerans* transmission.

**Fig. 3:**
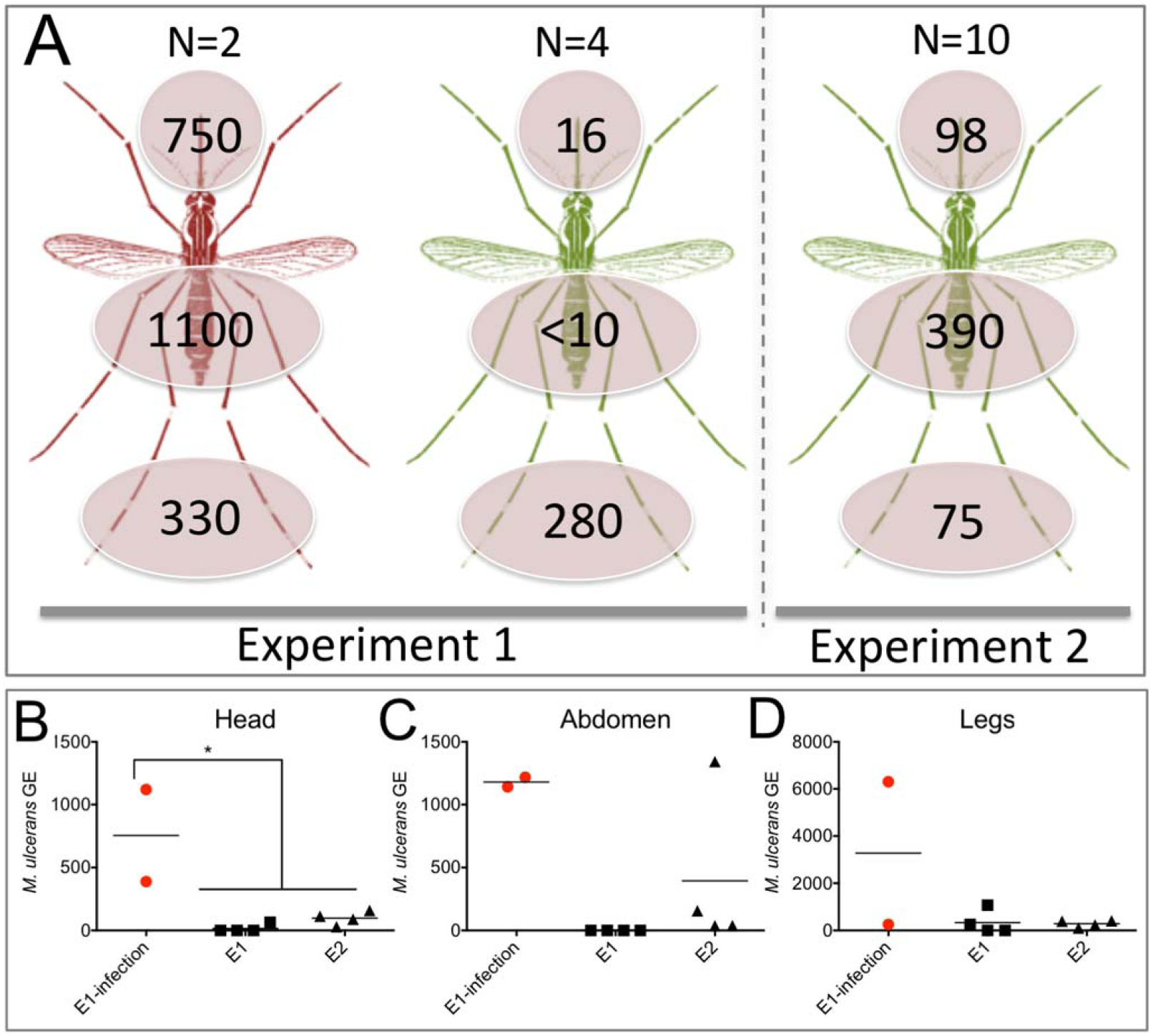
**Summary of *M. ulcerans* burden on mosquitoes post-feeding under two models of transmission.** (A) Visualization of the mean number of *M. ulcerans* detected per dissected mosquito segment, as assessed by *IS2404* qPCR and expressed as genome equivalents (GE). ‘N’ indicates the total number of mosquitoes tested. Red-shaded mosquitoes transmitted *M. ulcerans,* leading to mouse tail lesions. Green-shaded mosquitoes blood-fed on mouse-tails but lesions did not develop. (B, C, D) Plots of the individual qPCR results for each mosquito segment, listed by experiment. Red dots correspond to qPCR bacterial load for mosquitoes that transmitted *M. ulcerans* infection. Null hypothesis (no difference in bacterial load) was rejected (p<0.05)* (unpaired, two-tailed *t* test). Horizontal bar indicates the mean bacterial load per mosquito. The y-axis is GE and x-axis is experiment. The qPCR data for individual insects is contained in Supplementary Table S1.

### Estimation of incubation period and infectious dose of transmission model-1

Based on the time until a tail lesion was first observed, we estimated a median incubation period (IP) of 12 weeks (Fig. 4A). This result overlaps with the IP in humans for BU, estimated in different epidemiological studies from 4-10 weeks in Uganda during the 1960s [14] and 4 - 37 weeks in south east Australia [25]. We also estimated the infectious dose50 (ID50). We used six different concentrations of *M. ulcerans* to coat the tails of mice (n=5 mice/dilution) and observed the number of mice for each dilution that developed Buruli ulcer, allowing an ID_50_ estimate of 4 CFU (Fig. 4B). To our knowledge this is the first estimate of an *M. ulcerans* infectious dose and indicates that a surprisingly small quantity of this slow growing mycobacterium inoculated just below the skin is sufficient to cause disease.

**Fig. 4:**
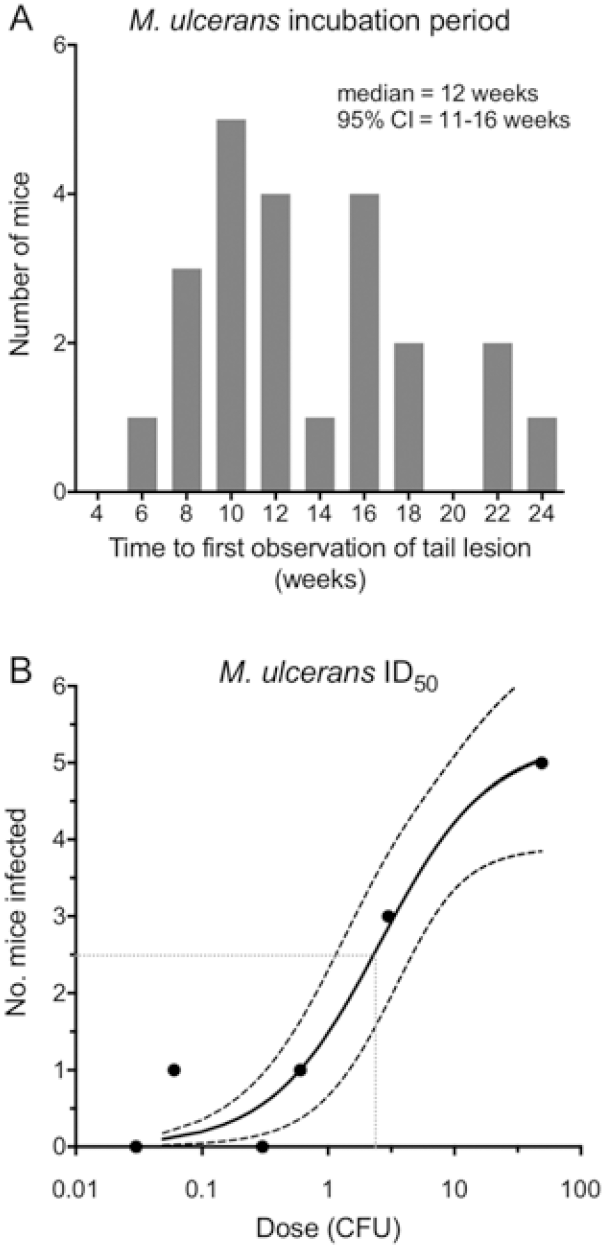
***M. ulcerans* incubation period and infectious dose_50_.** (A) Incubation period of *M. ulcerans* based on the time between sterile-needle puncture of an *M. ulcerans* contaminated mouse-tail and first observation of a lesion. (B) Estimated *M. ulcerans* ID_50_ for transmission model-1.

## Discussion

Here, we show for the first time a highly efficient atypical mode of mechanical transmission of *Mycobacterium ulcerans* to a mammalian host that implicates both biting insects and puncturing injuries. This research was designed around established frameworks for implicating vectors in disease transmission and provides the necessary causational evidence to help resolve the 80-year mystery on how *M. ulcerans* is spread to people [15, 31]. The efficient establishment of BU we have shown here via minor penetrating trauma through a contaminated skin surface is an atypical form of mechanical transmission *sensu lato (s.l.*) but it nonetheless satisfactorily fulfills one of the Barnett Criteria [31]. In vector ecology, mechanical transmission, *sensu strictu* (s.s.), is defined as a non-circulative process involving accidental transport of the pathogen. That is, the pathogen, in some fashion, nonspecifically associates or contaminates the mouthparts (stylet) of an arthropod vector. Insect mechanical transmission *s.s*. of BU implies that if *M. ulcerans* were ingested and then egested via regurgitation or salivation, the mechanism would act more like a syringe than a needle [32]. Such a mode of *M. ulcerans* disease transmission is supported by previous laboratory studies in which *Naucoris* and Belostmatid water bugs were contaminated via feeding on maggot prey that had been injected with *M. ulcerans* or fed naturally on dietary contaminated larval mosquito prey [23, 33].

Our demonstration in the current study of mechanical transmission *s.l* implies there are potentially multiple or parallel pathways of *M. ulcerans* infection [31]. Examples of bacterial diseases with multiple transmission modes include tularemia, plague and trachoma [34, 35]. Support for our mechanical transmission *s.l*. model also comes from the many field reports over the decades of *M. ulcerans* infection following trauma to the skin. Case reports have noted BU following a suite of penetrating injuries ranging from insect bites (ants, scorpions), snake bite, human bite, splinters, gunshot, hypodermic injections of medication and vaccinations [36–38]. Epidemiologists in Uganda during the 1960s and 70s suggested sharp-edged grasses might introduce the bacteria [39]. However, a recent laboratory study established that abrasions of the skin in Guinea pig models and subsequent application of *M. ulcerans* was not enough to cause an ulcer, however, this same study established that a subcutaneous injection would cause an ulcer [40]. As a sequel to this study in Guinea pigs, we raised the question of how likely it was that human skin could be sufficiently coated in *M. ulcerans* that an injury from natural or anthropogenic sources could lead to infection. Other explanations for the transmission of *M. ulcerans* include linkages with human behavior that increase direct contact with human skin and contaminated water [15]. A recent study from Cameroon recorded the persistence of *M. ulcerans* over a 24-month period in a waterhole used by villagers (including BU patients) for bathing [41]. A scenario could be envisaged where a villager’s skin surface becomes contaminated after bathing in such a water body and is primed for infection if (i) the concentration of bacteria is sufficiently high, and (ii) an inoculating event occurs. Whereas, in Australia, earlier studies have shown that *M. ulcerans* contamination of possum feces in and around the gardens of BU patients might present a similar skin surface contamination model in this region [17, 42]. Future experiments will address the possibility that insect vectors may be able to move *M. ulcerans* from one source and inject it into an animal or human.

Our focus on mechanical mosquito transmission *s.s*. arose from previous surveys in southeastern Australia where a strong association between *M. ulcerans* positive mosquitoes and human cases of BU has shown that *M. ulcerans* has not only been found on adult mosquitoes from both lab and field studies but also a biological gradient, where maximum likelihood estimates (MLE) of the proportion of *M. ulcerans*–positive mosquitoes increased as the number of cases of BU increased [19, 33, 43–46]. However, a recent study in Benin, West Africa found no evidence of *M. ulcerans* in association with adult mosquitoes [47]. The authors concluded that the mode of transmission might differ between southeastern Australia and Africa. Although, laboratory and fieldwork in West Africa suggest that aquatic insects, including mosquito larvae, play a role as reservoirs in nature for *M. ulcerans* that may be indirectly tied to transmission by serving as dispersal mechanisms [18, 23, 48]. Epidemiological studies have shown that direct contact with water is not a universal risk factor for BU [8, 11]. Prior exposure to insect bites and gardening are also independent risk factors for developing BU, while use of insect repellent is protective [11, 49].

Laboratory support to show mosquitoes can be competent vectors to spread BU is important additional evidence required to satisfy accepted vector ecology criteria for implicating insects in disease transmission [15, 31]. We found that infection was established following very minor penetrating trauma. *Aedes notoscriptus* mosquitoes feed by insertion of a stylet, sheathed within the proboscis, beneath the skin of a host. The stylet has a diameter around 10 *μ* M tapering to 1 *μ* M at its tip and extending 1-2 mm below the skin surface. We estimated the density of *M. ulcerans* on the mouse-tails surface was 100-200 CFU/mm^2^. Thus, the number of bacteria potentially injected during mosquito feeding through this contaminated surface is likely to be low, but this is consistent with our infectious dose estimates from needle-stick punctures, indicating an ID_50_ of only 4 CFU (Fig. 4B). There are strong parallels here with *Mycobacterium leprae*, the agent of leprosy. Like BU, the mode of transmission of the leprosy bacillus is unclear, but the infective dose is known to be very low (10 bacteria) and epidemiological evidence suggests multiple transmission pathways, including entry of the bacteria after skin trauma [50, 51]. Our infective dose estimate for *M. ulcerans* is consistent with observations that pathogens producing locally acting molecules to cause disease (e.g. the polyketide toxin mycolactone of *M. ulcerans*) have lower infective doses [52].

In summary, we have uncovered a highly efficient and biologically plausible atypical transmission mode of *M. ulcerans* infection via natural or anthropogenic skin punctures. Reduction of exposure to insect bites, access to clean water for bathing, and prompt treatment of existing BU are concrete measures likely to interrupt BU transmission.

## Supporting information Captions

**Supplementary Table S1:** IS2404 mosquito qPCR results

